# Genome-wide discovery of epistatic loci affecting antibiotic resistance using evolutionary couplings

**DOI:** 10.1101/325993

**Authors:** Benjamin Schubert, Rohan Maddamsetti, Jackson Nyman, Maha R. Farhat, Debora S. Marks

**Author notes:** Joint first author (Benjamin Schubert, Rohan Maddamsetti).

## Abstract

The analysis of whole genome sequencing data should, in theory, allow the discovery of interdependent loci that cause antibiotic resistance. In practice, however, identifying this epistasis remains a challenge as the vast number of possible interactions erodes statistical power. To solve this problem, we extend a method that has been successfully used to identify epistatic residues in proteins to infer genomic loci that are strongly coupled and associated with antibiotic resistance. Our method reduces the number of tests required for an epistatic genome-wide association study and increases the likelihood of identifying causal epistasis. We discovered 38 loci and 250 epistatic pairs that influence the dose needed to inhibit growth for five different antibiotics in 1,102 isolates of *Neisseria gonorrhoeae* that were confirmed in an independent dataset of 495 isolates. Many known resistance-affecting loci were recovered; however, the majority of loci occurred in unreported genes, including *murE* which was associated with cefixime. About half of the novel epistasis we report involved at least one locus previously associated with antibiotic resistance, including interactions between *gyrA* and *parC* associated with ciprofloxacin. Still, many combinations involved unreported loci and genes. Our work provides a systematic identification of epistasis pairs affecting antibiotic resistance in *N. gonorrhoeae* and a generalizable method for epistatic genome-wide association studies.

---

*Neisseria gonorrhoeae* is rapidly evolving to be untreatable. Clinical *N. gonorrhoeae* isolates already exhibit mono-resistance to extended-spectrum cephalosporins such as cefixime and ceftriaxone^1^”^4^ and other relevant antimicrobials^5^. More alarmingly, a new strain of *Neisseria gonorrhoeae* that is multi-resistant to the currently recommended therapy — a combination of azithromycin and ceftriaxone — was reported in the United Kingdom in March 2018^6^.

Epistasis can both enable and prevent the evolution of antibiotic-resistant bacteria^7^ such as *N. gonorrhoeae.* On the one hand, mutations that compensate for costly resistance variants^8^ can enable the evolution of multi-resistant bacteria. On the other hand, epistasis can constrain the spread of antibiotic resistance if the fitness cost of resistance prohibits its transmission or spread by horizontal gene transfer. Statistically, it is difficult to discover epistatic interactions that affect antibiotic resistance because the number of possible interactions to test is prohibitively large. To circumvent this problem, researchers have used prior knowledge such as known protein-protein interactions or network information to find candidate pairs for epistatic testing^9, 10^. Alternatively, others have restricted the statistical analysis to loci that individually are strongly associated with the trait of interest^11, 12^. In the end, neither of these approaches allow a systematic search for epistatic interactions.

To conduct a genome-wide association study^13^ (GWAS) that tests for epistatic associations in the face of the combinatorial explosion of statistical tests, we exploit sequence information by computing evolutionary couplings^14^ to identify epistatic interactions. In recent years, evolutionary couplings methods have made breakthroughs in *ab initio* protein and RNA 3D structure^14–16^, protein complex^17^, and mutation effects prediction^18^. Now, these methods are being adopted for bacterial genome analysis^19^. Evolutionary couplings are able to separate causal interactions from indirect correlations^20^ including, to a large extent, global correlations caused by population structure and phylogeny^21, 22^ or under-sampling. We hypothesized that evolutionary couplings, inferred from populations of bacterial pathogens, might represent functional or mechanistic dependencies between loci that affect bacterial fitness. If true, evolutionary couplings would be able to find epistatic interactions affecting phenotypes, and thus could be used to filter pairs for epistatic GWAS in a principled manner suggesting an agnostic alternative to pathway-based filtering approaches^9, 10^.

In this work, we report single nucleotide polymorphisms (SNPs) and epistatic interactions that were significantly associated with changes in minimal inhibitory concentration (MIC) in an exploratory dataset and subsequently confirmed in a separate dataset. The structure of the epistatic associations reveals the genetic architecture that underlies antimicrobial sensitivity and resistance in *N. gonorrhoeae.* Thus, our work provides a foundation for experimentalists and clinicians seeking to understand how epistatic interactions between genes affect the rapid evolution of antibiotic resistance in clinical pathogens such as *N. gonorrhoeae.*

## RESULTS

### N. gonorrhoeae *isolate genomes and drug resistance*

We extracted whole genome sequences and MICs of five antibiotics (Penicillin: PEN, Tetracycline: TET, Cefixime: CFX, Ciprofloxacin: CIPRO, and Azithromycin: AZI) from clinical studies of *N. gonorrhoeae* infection (Supplementary Table 1). Genome-wide association studies will often have some kind of validation test, though not strictly required to make the initial association. We therefore used one dataset of 1,102 strains collected in the USA^23^ for the initial exploration and a second dataset comprised of 495 strains^24^ collected in Canada^25^(*n* = 246) and England^26^ (*n* = 249), which we used for independent confirmation (Supplementary Table 1). The phylogenetic analysis of all 1,597 isolates showed considerable diversity within both cohorts (Supplementary Figure 1, Methods). The cohorts nevertheless differed in lineage composition, supporting the use of the second dataset as confirmatory. In a phenotypic analysis of susceptibility to the antibiotics, we found that 92% of isolates from both cohorts were resistant to at least one drug, using clinically defined thresholds from the European Committee of Antimicrobial Susceptibility Testing^27^ (Figure 2). Moving forward to the analysis of resistance association, using *N. gonorrhoeae* FA1090 as a reference, we identified non-synonymous SNPs and SNPs in non-coding regions affecting gene expression^28^ that had a minor allele frequency ≥ 0.5%, resulting in 8,686 loci (Methods).

**Figure 1.**
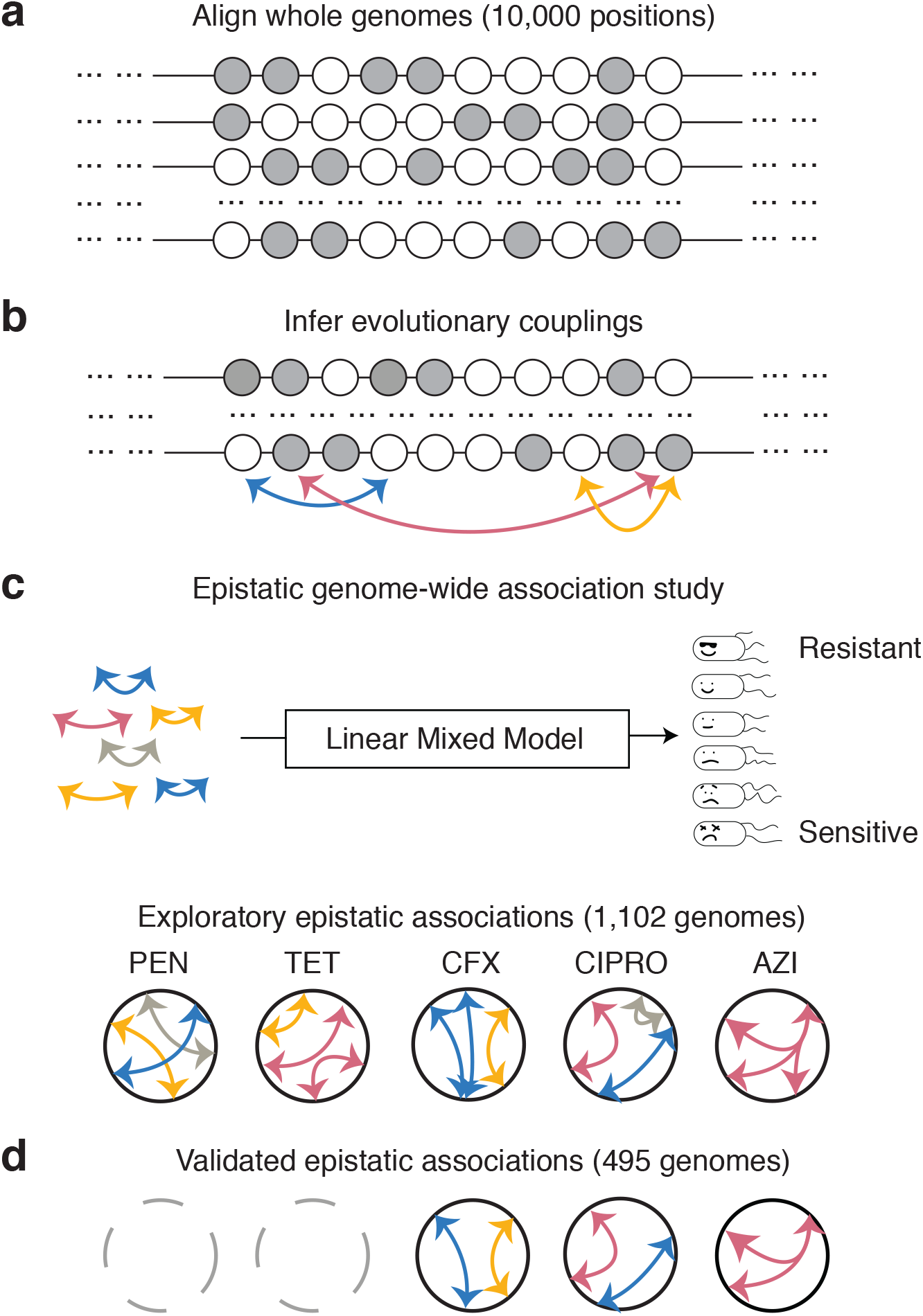
Evolutionary couplings power epistatic GWAS. **a**) First, non-synonymous SNPs and non-coding variants with experimentally-derived functional annotation are identified and used to generate a whole-genome alignment. **b**) Highly evolutionarily coupled loci are identified using an undirected graphical model given the multiple sequence alignment. **c**) The inferred evolutionary couplings are then tested in an exploratory set of 1,102 *Neisseria gonorrhoeae* strains, using a linear mixed model, for their epistatic association to minimum inhibitory concentrations for five antibiotics: penicillin (PEN), tetracycline (TET), cefixime (CFX), ciprofloxacin (CIPRO), and azithromycin (AZI). **d**) Significant epistatic associations are then validated in an independent and geographically distinct set of 495 *N. gonorrhoeae* genomes.

**Figure 2.**
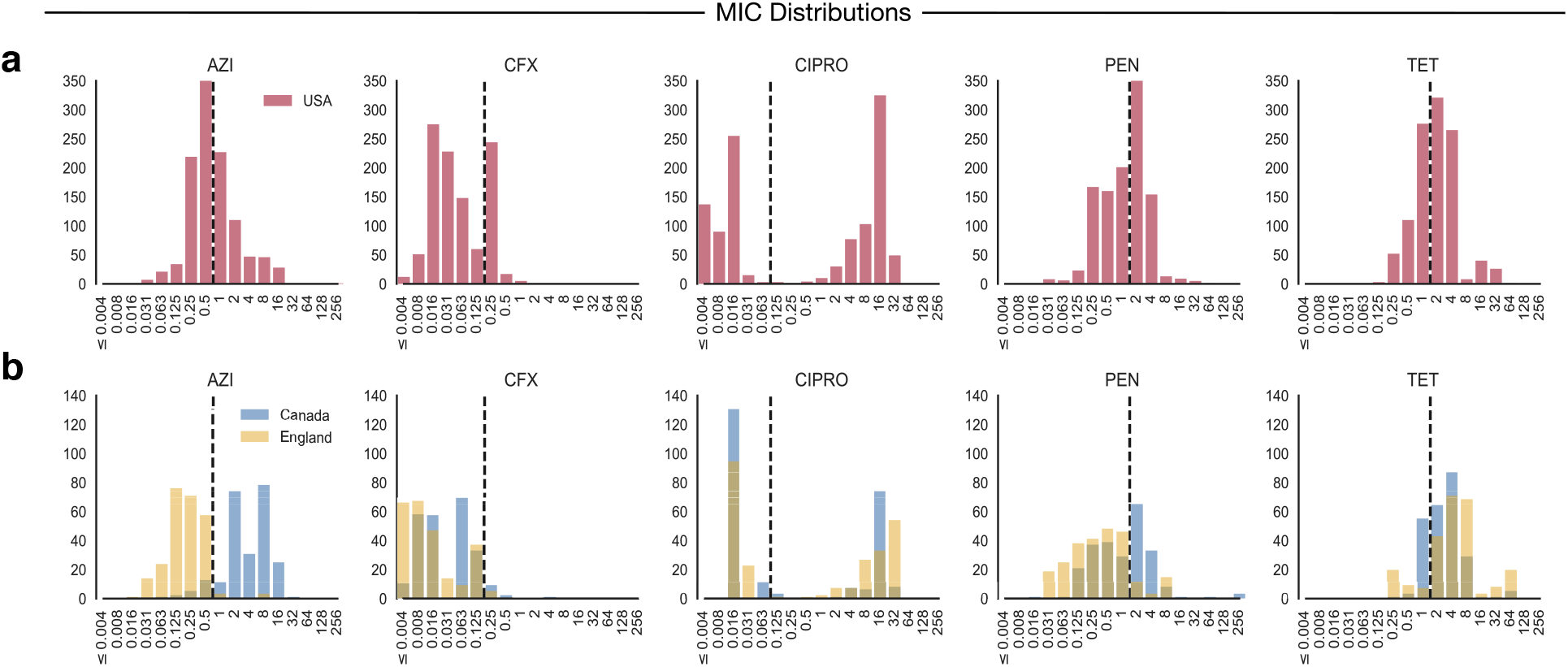
Distributions of minimum inhibitory concentration (MIC), in milligrams per liter, for the *N. gonorrhoeae* strains studied. The dotted lines indicate clinical breakpoints for *N. gonorrhoeae* as defined by the European committee on antimicrobial susceptibility testing (EUCAST). **a**) The MIC distribution of the 1,102 strains in the exploratory test dataset. In the test set, 537 strains were resistant to penicillin (PEN; resistance threshold > 1 mg/L), 661 were resistant to tetracycline (TET; > 1 mg/L); 266 were resistant to cefixime (CFX; > 0.125 mg/L); 602 were resistant to ciprofloxacin (CIPRO; > 0.06 mg/L); and 459 were resistant to azithromycin (AZI; > 0.5 mg/L). b) The MIC distribution of the 495 strains in the confirmatory dataset. The confirmatory set contained 141 PEN, 400 TET, 17 CFX, 231 CIPRO, and 231 AZI resistant.

### Probabilistic model to capture genome-wide interactions

To test for epistatic contributions to the observed antibiotic resistance, we would have to test over 37 million combinations of the 8,686 loci— resulting in low statistical power and a high probability of identifying spuriously correlated pairs. Since it seems reasonable to assume that only a small number of pairs of loci, if any, are causally related to the resistance phenotype, a logical approach would be to simply test the most correlated pairs across the samples. However, many of those pairs are likely to be non-causal, due to population structure in the data that results in transitive correlations. This problem is seen in many areas of biological data analysis: correlation does not imply causation^20, 29^. Here we solve this by applying a maximum entropy model to identify which pairs of loci best explain all other observed pairs in the data. The model and inference approaches we develop are based on a method that identifies causal dependencies and epistasis between residues in proteins and RNAs that has led to successful folding of 3D structure^14,20,30^ and prediction of mutation effects^18^ from sequence alone.

We therefore computed the epistatic relationships between the 8,686 loci before exploring the association to antibiotic resistance phenotypes. Our statistical model associates the genome sequence *σ* with a probability distribution *P(σ)* at equilibrium as

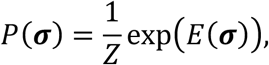

where Z is a normalization constant. We define *E(**σ**)* as the sum of coupling terms ***J**_ij_* between every pair of loci in a sequence and a locus-wise bias term ***h**_i_*:

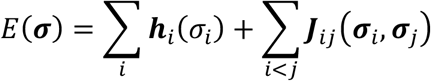

We use regularized pseudolikelihood maximization inference^31^ to compute the parameters ***J**_ij_* and ***h**_i_*. To measure the evolutionary coupling (EC) strength between pairs of loci, the inferred parameters ***J**_ij_* are summarized using the Frobenius norm^31^:

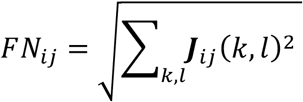

To calculate the final EC score, we corrected the FN scores for population-structure and under-sampling using the average product correction^32^. From the full set of all pairs (~ 37 million) we selected the subset that had a probability of more than 95% for being in the signal distribution when fitting the coupling strengths to a two-component mixture model (Methods). This resulted in a high-confidence set of 242,360 pairs involving 7,868 loci in 1,438 genes with the majority in protein–coding genes (7,437); 178 within transcription start sites or 5’ untranslated regions; 224 within promoters; and 29 within ribosomal RNAs (Supplementary Table 2).

### Genome-wide single-site analysis recapitulates known associations and predicts new ones

Before testing the epistatic pairs of loci, we performed a single-site GWAS analysis. We used a linear mixed model to measure the association strength of individual loci with the measured change in minimum inhibitory concentrations, correcting for population structure, as in standard GWAS approaches^33, 34^ (Methods). We tested all 8,686 loci for single association, controlling for multiple hypothesis testing using a Bonferroni correction with an adjusted *α* set to 0.05. Nearly half (38/82) of the significant associations in the exploratory dataset were also significant in the confirmatory dataset (Tables 1 and 2, Supplementary Table 3).

**Table 1.** Single loci associated with change in sensitivity or resistance to Ciprofloxacin (CIPRO)

| Locus | Gene | Variant compared to FA1090 | *p*-value | $\beta$ | Count | Minor allele frequency in resistant isolates |
| --- | --- | --- | --- | --- | --- | --- |
| **620917\*** | ***gyrA* (NGO0629)** | **S91F** | **$8.25 \times 10^{-101}$** | **2.727** | **236** | **230/231** |
| **620905\*** | ***gyrA* (NGO0629)** | **D95(A/G)** | **$1.98 \times 10^{-43}$** | **2.091** | **24/202** | **222/231** |
| **1210781\*** | ***parC* (NGO1259)** | **S87R** | **$9.96 \times 10^{-18}$** | **1.929** | **117** | **173/231** |
| 894348* | *dldH* (NGO0915) | A88V† | $6.68 \times 10^{-9}$ | -1.265 | 286 | 14/231 |
| 1167114* | *gshB* (NGO1217) | E221K | $2.71 \times 10^{-8}$ | 0.920 | 205 | 194/231 |
| 1687733* | *comF* (NGO1726) | F43Y | $1.86 \times 10^{-7}$ | -1.004 | 127 | 19/231 |
| 93476 | NGO0086 | D363E | $1.66 \times 10^{-5}$ | -0.885 | 198 | 12/231 |
| 1167527 | *gshB* (NGO1217) | E83A† | $1.91 \times 10^{-5}$ | 0.647 | 255 | 195/231 |
| 89733 | *pglD* (NGO0083) | A360T | $2.07 \times 10^{-5}$ | 0.728 | 171 | 162/231 |
| 1393066* | *cirA* (NGO1430) | R122P | $2.83 \times 10^{-5}$ | 0.947 | 155 | 147/231 |
| 1921503* | *doxX* (NGO1948) | L139R | $4.09 \times 10^{-5}$ | 0.837 | 219 | 184/231 |
| 1923171* | *prfB* (NGO1951) | -84 A→G | $4.93 \times 10^{-5}$ | -0.930 | N/A | 17/231 |
| 447535 | *fimT* (NGO0452) | G100D† | $9.02 \times 10^{-5}$ | -0.553 | 270 | 22/231 |
| 1162600 | *pyrG* (NGO1212) | -12 G→A | 0.000176 | 0.647 | N/A | 172/231 |
| 1162602 | *pyrG* (NGO1212) | -14 A→G | 0.000176 | 0.647 | N/A | 172/231 |
| 964051* | NGO0993 | K161E† | 0.000389 | -0.526 | 351 | 5/231 |
| 1378462 | *nqrB* (NGO1414) | A25V | 0.000661 | -0.578 | 138 | 13/231 |
| 565114 | *uvrC* (NGO0578) | N258D | 0.000696 | 0.664 | 227 | 182/231 |
| 1747633 | *gapA-B* NGO1776 | V267A† | 0.000882 | -0.568 | 293 | 20/231 |
| 1373003 | NGO1407 | L66M | 0.00116 | -0.666 | 116 | 1/231 |
| 1372748 | NGO1407 | -61 A→G | 0.00116 | -0.666 | N/A | 1/231 |
| 1373005 | NGO1407 | L66M | 0.00116 | -0.666 | 116 | 1/231 |

NOTE— Loci in bold are known resistance variants. Asterisk (\*) loci are associated with significant epistatic interactions, shown in Figure 3. Coding variants have the minor allele on the right († minor allele on left). Non-coding variants are labeled with the distance upstream (–) from the closest gene. $\beta$ is the regression coefficient associated with the minor allele variant in the linear mixed model, run on the confirmatory dataset. Reported *p*-values follow a Bonferroni correction for multiple tests. The count of each variant (with respect to FA1090) in *N. gonorrhoeae* isolates from the confirmatory dataset is reported, as is the fraction of resistant isolates in the confirmatory dataset with the minor allele variant.

**Table 2.** Single loci associated with change in sensitivity or resistance to Azithromycin (AZI), Penicillin (PEN), Tetracycline (TET), and Cefixime (CFX)

| Antibiotic | Locus | Gene | Variant | *p*-value | $\beta$ | Count | Minor allele frequency in resistant isolates |
| --- | --- | --- | --- | --- | --- | --- | --- |
| **AZI** | **1116541\*** | **23S rRNA (NGOr02)** | **C2617T (C2611T)** | **2.19x10<sup>-31</sup>** | **0.889** | **N/A** | **143/231** |
| **PEN** | **1789058** | ***porB* (NGO1812)** | **A121(D/G)** | **2.45x10<sup>-4</sup>** | **0.348** | **123/7** | **69/141** |
| PEN | 1789148 | *porB* (NGO1812) | V151A | 4.78x10<sup>-3</sup> | -0.249 | 178 | 6/141 |
| TET | 1789088 | *porB* (NGO1812) | F131Y | 1.33x10<sup>-3</sup> | -0.174 | 135 | 83/400 |
| CFX | 1516685* | *penA* (NGO1542) | V160A | 2.14x10<sup>-15</sup> | 0.958 | 96 | 16/17 |
| CFX | 1516646* | *penA* (NGO1542) | N173S | 2.14x10<sup>-15</sup> | 0.958 | 96 | 16/17 |
| **CFX** | **1515528\*** | ***penA* (NGO1542)** | **G546S** | **5.08x10<sup>-12</sup>** | **0.859** | **95** | **15/17** |
| CFX | 1516861* | *penA* (NGO1542) | D101E | 1.54x10<sup>-10</sup> | 0.954 | 103 | 16/17 |
| CFX | 1516798* | *penA* (NGO1542) | V122(F/V) | 1.54x10<sup>-10</sup> | 0.954 | 103 | 16/17 |
| CFX | 1514399 | *murE* (NGO1541) | A331(E/G) | 1.41x10<sup>-9</sup> | 0.835 | 103 | 14/17 |
| CFX | 1514450 | *murE* (NGO1541) | A314G | 1.41x10<sup>-9</sup> | 0.835 | 103 | 14/17 |
| CFX | 1516552* | *penA* (NGO1542) | D204E | 7.54x10<sup>-9</sup> | 0.802 | 114 | 16/17 |
| CFX | 1516556* | *penA* (NGO1542) | E203G | 7.54x10<sup>-9</sup> | 0.802 | 114 | 16/17 |
| CFX | 1516563* | *penA* (NGO1542) | Y201H | 7.54x10<sup>-9</sup> | 0.802 | 114 | 16/17 |
| CFX | 1515252 | *murE* (NGO1541) | A47T† | 2.83x10<sup>-4</sup> | 0.465 | 344 | 16/17 |
| CFX | 1515263 | *murE* (NGO1541) | P43(L/Q)† | 2.83x10<sup>-4</sup> | 0.465 | 4/344 | 16/17 |

NOTE– See Table 1 Note for explanation.

#### Ciprofloxacin

Of these 38 loci, 22 were associated with CIPRO with known loci in *gyrA* (2 loci) and*parC* as the most significant (p= 8.25×10^-101,^ 1.98×10^-43^ and 9.96×10^-18^ respectively) (Table 1). We also observed significant associated loci in 20 genes and four non-coding regions (Table 1). The two strongest novel associations, found in *dldH* (dihydrolipoyl dehydrogenase, p.A88V, *p* = 6.68×10^-9^) and *gshB* (glutathione synthetase, p.E221K, *p =* 2.71×10^-8^) are involved in oxidative stress response^35^ which can be induced by fluoroquinolones^36^ (e.g. ciprofloxacin). The *dldH* variant A88V is on the homodimer interface of the multimeric enzyme, suggesting that multimerization stability is part of its mechanism (Figure 3). Our analysis suggests that the *dldH* minor allele variant (alanine in the validation dataset) causes greater CIPRO sensitivity, since it is uncommon in resistant isolates (14/231) and its regression coefficient (β) is negative. By contrast, the minor allele variant E221K in *gshB* is associated with a higher CIPRO MIC and a positive regression coefficient (β), suggesting that the lysine variant increases resistance. The highest density of CIPRO associations was found in *gyrA;* the AsnC family transcriptional regulator NGO1407 (three loci); *gshB;* and in the 5’ untranslated region of *pyrG* (two loci each).

**Figure 3.**
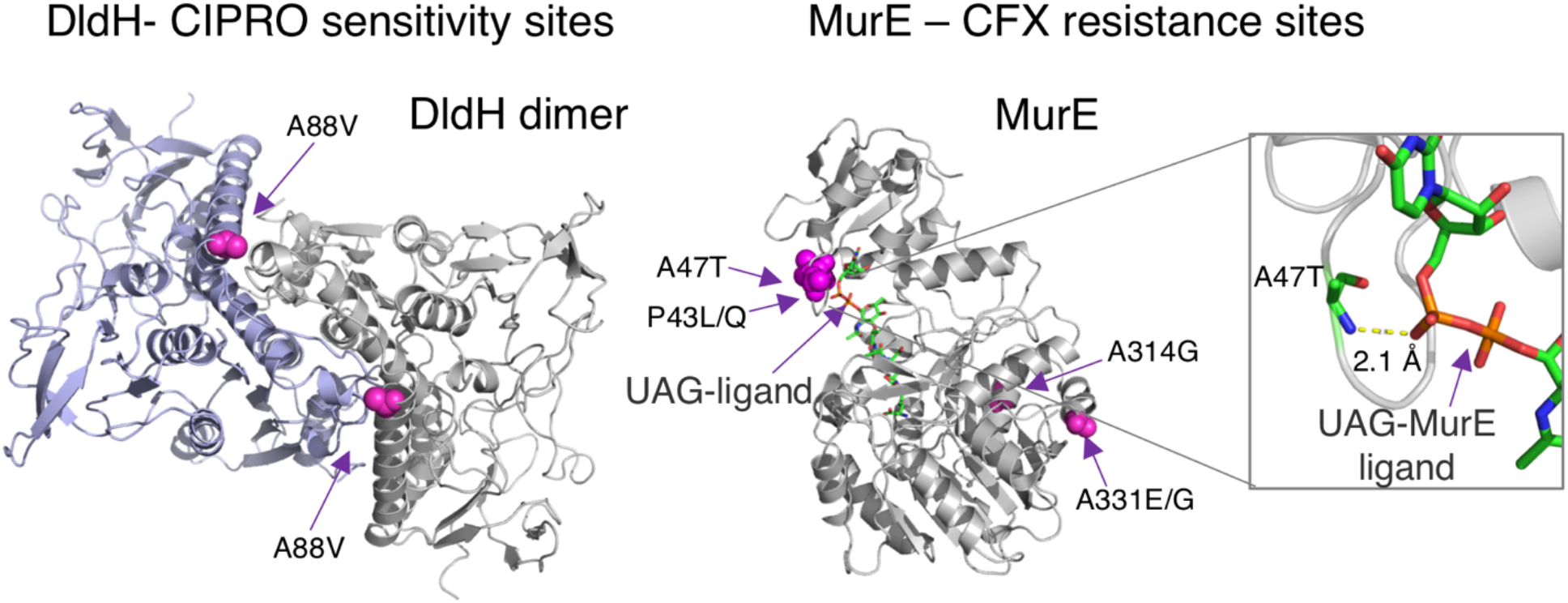
Novel associations that occur in structural binding sites. **Left**: *dldH* A88 (magenta), associated with CIPRO resistance, maps onto the interface of the homodimer shown in the 3D structure of the homologous protein in *Pisum sativum,* 1dxl^47^. **Right**: Two out of four variants associated with CFX resistance in *murE* map onto the enzyme active site, as shown in the 3D structure of the homologous protein in *E.coli,* 1e8c^48^, co-crystalized with its substrate uri-dine-5’s-diphosphate-n-acetylemuramoyl-l-alanine-d-glutamate (UAG). See Tables 1 and 2 for details.

#### Cefixime

Eight of our CFX-associated loci are in the antibiotic target *penA* (PBP2, penicillin-binding protein 2). *penA* is mosaic in the test isolates^23^ and only one of these eight associations has been shown to affect resistance^4^ (p.G546S, *p* = 5.08×10^-12^). The remaining four significantly associated loci are in the cell-wall biosynthesis gene, *murE.* Mutations in *murE* increase β-lactam resistance in *S. pneumoniae*^37^, but to our knowledge, have not been reported in *N. gonorrhoeae* (Figure 3).

#### Penicillin, Tetracycline, and Azithromycin

We recover one of the known*porB* sites associated with increased resistance^23^ (A121D/G/V, *p* = 2.45×10^-4^) in our confirmation dataset. We also identified previously unreported loci in *porB* that are likely to be associated with increased antibiotic sensitivity (its regression coefficient β is negative). One was associated with PEN (p.V151A, *p* = 4.73×10^-3^) and another was associated with TET (p.F131Y, *p* = 1.33×10 ^3^). Since variants in *porB* have also been associated with antibiotic resistance, this shows that *porB* variation can increase sensitivity as well as resistance to antibiotics. The only single association we see for AZI is the previously known locus in 23S rRNA g.C2617T, previously described as C2611T *(p* = 2.19×10^-31^).

### Genome-wide association identifies epistatic pairs associated with antibiotic resistance

Next, we tested all 242,360 pairs of loci with high-scoring evolutionary couplings for epistatic association. To assess the contribution of each epistatic effect, we compared the full interaction model against a model only containing the individual sites as additive effects. As before, we applied a Bonferroni correction for multiple testing at an adjusted *α* = 0.05.

We verified 250 epistatic associations in the confirmatory dataset (Figure 4, Table 3), out of 729 significant epistatic associations in the exploratory dataset (Supplementary Tables 4 and 5). We confirmed epistatic interactions to CIPRO, CFX and AZI but not to TET or PEN.

**Table 3.** Selected set of epistatic pairs of loci associated with change in antibiotic sensitivity or resistance (Detailed results in Supplementary Table S5)

| Antibiotic | Locus i | Locus j | Gene i | Gene j | Variant i | Frequency in resistant isolates | Variant j | Minor allele frequency in resistant isolates | $\beta$ | p-value |
| --- | --- | --- | --- | --- | --- | --- | --- | --- | --- | --- |
| AZI | 1116541 | 1230597 | 23S rRNA | NGO1277 | C2617T | 315/690 | D173G | 358/690 | 0.877 | $5.77 \times 10^{-29}$ |
| AZI | 1116541 | 1427960 | 23S rRNA | *opA54* (NGO1463a) | C2617T | 315/690 | V123L | 32/690 | 0.877 | $5.77 \times 10^{-29}$ |
| AZI | 1116541 | 1530961 | 23S rRNA | *putA* (NGO1552a) | C2617T | 315/690 | E1169G | 377/690 | 0.877 | $5.77 \times 10^{-29}$ |
| AZI | 1116541 | 1789096 | 23S rRNA | *porB* (NGO1812) | C2617T | 315/690 | N134D/H | 352/690 | 0.877 | $5.77 \times 10^{-29}$ |
| CFX | 1516646 | 1517334 | *penA* (NGO1542) | *ftsL* (NGO1543) | N173S | 277/283 | L52V | 0/283 | 0.956 | $1.63 \times 10^{-10}$ |
| CFX | 1516646 | 2025976 | *penA* (NGO1542) | *efeO* (NGO2050) | N173S | 277/283 | S360G | 0/283 | 0.959 | $1.63 \times 10^{-10}$ |
| CFX | 1516685 | 2070085 | *penA* (NGO1542) | *fetA* (NGO2093) | V160A | 277/283 | A638V | 132/283 | 0.959 | $1.63 \times 10^{-10}$ |
| CFX | 1516685 | 1665722 | *penA* (NGO1542) | *yebE* (NGO1709) | V160A | 277/283 | K90E | 258/283 | 0.959 | $1.63 \times 10^{-10}$ |
| CIPRO | 620905 | 1210781 | *gyrA* (NGO0629) | *parC* (NGO1259) | D95A/G | 270/833 | S87R | 702/833 | 1.984 | $1.26 \times 10^{-30}$ |
| CIPRO | 620917 | 1210781 | *gyrA* (NGO0629) | *parC* (NGO1259) | S91F | 239/833 | S87R | 702/833 | 2.413 | $1.73 \times 10^{-45}$ |
| CIPRO | 620917 | 2028211 | *gyrA* (NGO0629) | *glmU* (NGO2053) | S91F | 239/833 | T425A | 191/833 | 2.413 | $1.73 \times 10^{-45}$ |
| CIPRO | 620917 | 1974020 | *gyrA* (NGO0629) | *birA* (NGO2001) | S91F | 239/833 | E261K | 56/833 | 2.413 | $1.73 \times 10^{-45}$ |
| CIPRO | 620917 | 693931 | *gyrA* (NGO0629) | *hsdS* (NGO0699) | S91F | 239/833 | A173T | 47/833 | 2.413 | $1.73 \times 10^{-45}$ |
| CIPRO | 620917 | 894348 | *gyrA* (NGO0629) | *dldH* (NGO0915) | S91F | 239/833 | A88V | 46/833 | 2.413 | $1.73 \times 10^{-45}$ |
| CIPRO | 620917 | 1167114 | *gyrA* (NGO0629) | *gshB* (NGO1217) | S91F | 239/833 | E221K | 242/833 | 2.413 | $1.73 \times 10^{-45}$ |
| CIPRO | 620917 | 1522835 | *gyrA* (NGO0629) | *ftsN* (NGO1549) | S91F | 239/833 | Q142R | 244/833 | 2.413 | $1.73 \times 10^{-45}$ |

NOTE— Loci in bold are known resistance variants. Non-coding variants are labeled with the distance upstream (–) from the closest gene. $\beta$ is the regression coefficient associated with each locus in the linear mixed model. Reported *p*-values follow a Bonferroni correction for multiple tests. The count of each variant in all *N. gonorrhoeae* isolates (both test and confirmatory datasets) is reported, as is the fraction of resistant isolates (both test and confirmatory datasets) with that variant.

**Figure 4.**
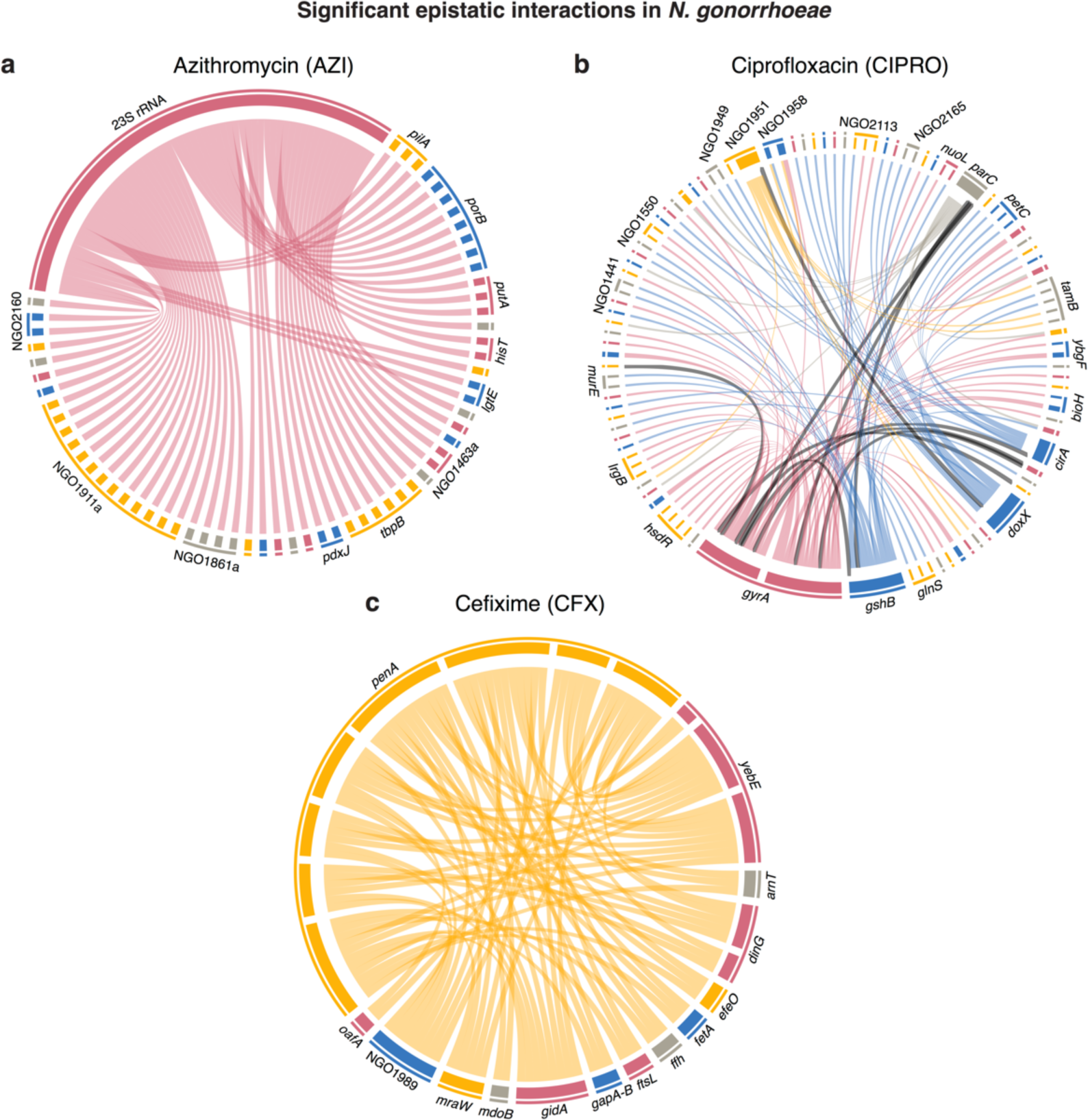
Significant epistatic interactions affecting antibiotic resistance and sensitivity in *Neisseria gonorrhoeae.* Edges between pairs of positions that are both individually associated with changes in minimum inhibitory concentration (MIC) are outlined in black, **a**) Epistatic interactions affecting resistance to azithromycin (AZI). All interactions are connected to a single C2617T variant in 23 S rRNA, which is known to cause significant resistance to AZI. **b**) Epistatic interactions affecting resistance to ciprofloxacin (CIPRO), **c**) Epistatic interactions affecting resistance to cefìxime (CFX).

All 250 epistatic interactions involved loci that were confirmed in our single locus GWAS. Ten epistatic associations connected two loci which were each significant in the single locus GWAS. The remaining 240 interactions connected an associated locus to a non-associated locus. In addition, most of the known resistance variants that we identified in our single locus GWAS (5/6) also occurred in our epistatic GWAS analysis (141 occurrences). One hundred and eighteen of the confirmed epistatic associations were associated with CIPRO MIC change; 69 with CFX; and 63 with AZI. Although no interactions had both loci of the pair shared across antibiotics, four genes had epistatic associations with multiple antibiotics, including *rmsE,* a 16S rRNA (uracil(1498)-N(3))-methyltransferase between shared between AZI and CIPRO.

#### Ciprofloxacin

Epistatic interactions associated with CIPRO involved 79 genes (Figure 3B), and the top 15 most significant (all at *p* = 1.73×10^-45^) involved the well-known S91F resistance variant in *gyrA*^5^. Two of the strongest epistatic pairs included interactions between the two known resistance-causing loci in *gyrA* and a resistance-causing locus in *parC (gyrA* p.S91F to *parC* p.S87F, *p* = 1.73×10^-45^; *gyrA* p.D95A/G to *parC* p.S87F, *p* = 1.26×10^-30^). The gene products of *parC* and *gyrA* are both topoisomerases type II proteins and these well-known loci, S91F (gyrA) and S87F (*parC*), are in homologous positions close to the DNA and ciprofloxacin binding sites where the resistance variants allow the enzyme to re-ligate DNA^38^ in the presence of an antibiotic. To our knowledge this is the first report of epistatic effects between these loci in *N. gonorrhoeae*; notably, homologous epistatic interactions (affecting both ciprofloxacin resistance and fitness) have been seen in experiments with *S. pneumoniae*^39^.

We also identified interactions between *gyrA* S91F and *birA* E261K, *(p* = 1.73×10^-45^) and *glmU* T425A *(p* = 1.73×10^-45^) respectively. The gene *birA* is known to increase sensitivity to ciprofloxacin when underexpressed in *Mycobacterium smegmatis^40^*, while GlmU’s protein abundance increases under ciprofloxacin stress in *Salmonella typhimurium,* possibly modulating cell wall permeability^41^. In total 15 genes contained three or more epistatic associations, dominated by inner and outer-membrane proteins *(doxX, lrgB, cirA, petC,* NGO1958, *tamB),* some of which are associated with oxidative stress^35^.

#### Azithromycin

All epistatic associations to AZI involved a single, known antibiotic resistance variant C2617T (identical to C2611T in previous reports^23^) in the 23S rRNA (Table 2 and Figure 3A). Of the eleven genes containing two or more associated loci, seven were either membrane proteins (NGO1496, NGO1861a, NGO1463a,porB), cell wall biosynthesis proteins *(lgtE)* or pilus proteins (NGO 1911a, *pilA).*

#### Cefixime

By contrast with CIPRO, epistatic associations with CFX involved only 15 genes, each with multiple associated loci. Eleven epistatic interactions involved the known single resistance locus (p.G546S) in *penA.* The fifteen genes were dominated by cell membrane, cell wall, and cell wall biosynthesis proteins *(penA*, *fetA, efeO, yebE, mdoB, ffh, arnT, oafA),* but also included cell division proteins (*ftsL*, *gidA*), and DNA repair (*dinG*).

Taken together, significantly associated epistatic interactions often involved known antibiotic resistance loci. Of the interactions that did not involve known antibiotic resistance loci, some were found in genes associated with cell division, and oxidative stress. The majority of novel epistatic loci, however, were found in genes encoding inner- and outer-membrane proteins, or cell wall biosynthesis proteins that potentially affect *N. gonorrhoeae’s* permeability to antibiotics^42–44^.

## DISCUSSION

Here, we report the results of the first systematic GWAS of loci and epistatic interactions affecting antimicrobial sensitivity and resistance in *N. gonorrhoeae.* As expected, we recover many known resistance-causing variants, and many of the epistatic interactions that we discovered involve known resistance-causing variants.

In our study, reported loci can cause either heightened *sensitivity* or greater *resistance* to antimicrobials. Examples for heightened sensitivity include loci in *dldH* and *comF* that are associated with CIPRO (Table 1). We also found several unreported variants and interactions that associate strongly with greater antibiotic resistance, including several variants in *murE* linked to CFX resistance and two interactions between *gyrA* and *parC* linked to CIPRO resistance. Each antibiotic showed distinct epistatic association patterns. While epistatic associations with CIPRO occurred in many genes, all epistatic associations with AZI involved a single resistance locus in 23S rRNA, and all epistatic associations with CFX involve loci in a few genes, mostly concentrated in the peptidoglycan synthesis gene *penA.* These observations suggest that the mechanism of CIPRO-induced death involves many genes across diverse cellular pathways, including oxidative stress, while the mechanism of CFX and AZI killing is focused on disabling peptidoglycan synthesis and 23S rRNA, respectively.

In principle, GWAS is able to identify causal variation. However, our study focuses only on point mutations found by comparison to the *N. gonorrhoeae* FA1090 reference genome. Thus, the effects of horizontally-transmitted genes and more complex variants such as gene duplications, indels, and mobile element transpositions will be missed. Some of these missing variants may be hidden yet causal variables that affect antibiotic resistance. If so, the effects of these missing variants will either be absent or will be associated to non-causative markers in our data that closely track the presence of the true, but missing, causal variants. We find some evidence of missing causal variables in our analysis as many epistatic associations are linked to genes that are highly variable in the genome alignment, such as transferrin-binding proteins^45^. Thus, it is possible that some associations involving highly variable genes are actually markers that represent the closest possible association to hidden causal variants. Nevertheless, our work demonstrates that evolutionary couplings combined with standard GWAS methods is a promising new approach for discovering epistatic interactions affecting antibiotic resistance in a multi-resistant pathogen, *N. gonorrhoeae.* The loci and interactions captured may support efforts in predicting antibiotic resistance from whole genome sequence data^46^. We anticipate that our findings as well as our method will be valuable in unraveling the genetics of complex traits in general.

## ACKNOWLEDGEMENTS

*We thank members of the Marks lab and Chris Sander for his support during this research project. The authors declare no conflict of interest*.

## AUTHOR CONTRIBUTIONS

D.S.M. conceived the project and supervised research; D.S.M, B.S and R.M designed and planned research; B.S. implemented the analysis methods; B.S. and R.M. analyzed data with the help of J.N., M.R.F., and D.S.M.; B.S., R.M, and D.S.M. wrote the paper. All authors reviewed and edited the manuscript.

## METHODS

### Alignment construction and annotation

We used *breseq*^1^ to call variants in each isolate with reference to the *Neisseria gonorrhoeae* FA1090 genome. To identify variants in ribosomal rRNAs, we masked all except the first copy of the rRNA operon in the reference genome. We ran *breseq* in consensus-mode using the masked reference genome. We filtered variants in the consensus-mode runs for single nucleotide polymorphisms (SNPs), then mapped the SNPs onto the reference sequence using *gdtools* (part of *breseq)* to generate a genome alignment. We ran *breseq* in polymorphism-mode to infer the copy number of rRNA variants in each isolate. The FA1090 reference genome was used to annotate the location of coding regions (CDS) and other genomic features. Promoters and 5’ untranslated regions associated with experimentally determined RNA transcripts^2^ were annotated as follows. Regions bounded by the transcriptional start site and the location of the first gene on the transcript were annotated as 5’ untranslated regions, as long as the annotated region was no more than 150 nucleotides (nt) long. This decision rule was based on experimental results on the distribution of 5’ untranslated region lengths in *Bacillus subtilis, Escherichia coli,* and *Pseudomonas aeruginosa^3^*. Sequences up to 70 nt upstream of the transcript (or, the end of the upstream gene on the same strand if within 70 nt) were annotated as promoters, following Remmele *et al.^2^.* The genome alignment was filtered for variants that occurred in at least 5 strains (0.5% minor allele frequency). Synonymous variants within CDS were excluded from the alignment.

### Population structure analysis

We used FastTree^4^ to make an approximately maximum-likelihood phylogeny using a genome alignment of all 1,597 strains.

### Genome-wide evolutionary couplings inference

To identify strongly coupled positions within a genome of length L, we fit a *q =* 2 state undirected graphical model of the form:

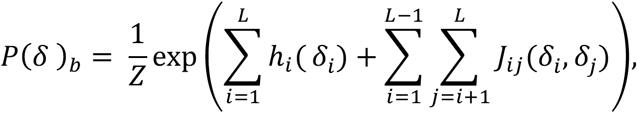

using a multiple sequence alignment of *N* bacterial genomes. Minor alleles were encoded as ‘1’ and major alleles were encoded as ‘0’. To prevent overfitting, we used L2-regularized pseudolikelihood estimation, similar to previous work^7–11^. The regularization parameters for the site-specific term was set to *λ_h_* = 0.01 and the pair-wise term was set to *λ_e_* = *λ_h_*(*L* – 1)*q*. To summarize the evolutionary couplings strength of each pair of loci, we use the Frobenius norm (FN)^7^:

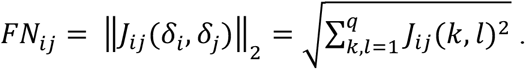

### Identification of highly evolutionarily coupled loci

To correct for undersampling and phylogenetic bias in the alignment, we adjusted the summarized FN-scores with an average-product correction (APC)^12^. Following Toth-Petroczy *et al*.^11^, a two-component mixture model was fitted to the corrected FN-score distribution to identify strongly evolutionarily coupled genomic loci:

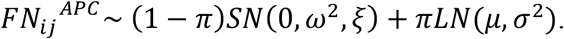

Here *π* represents the mixing parameter; SN is a skew-normal distribution with location at 0 and unknown scale *ω* and shape ξ parameters; and LN represents a log-normal distribution with unknown mean *μ* and variance *σ*. All evolutionary couplings (ECs) whose posterior probability of membership in the log-normal component *p_ij_* is larger than 0.95 are deemed significant and used for later analysis.

### Detection of antibiotic-resistant loci and co-evolutionarily coupled pairs

To test single-locus and epistatic associations to a specific phenotype, we employ a linear mixed model^13, 14^ of the following form:

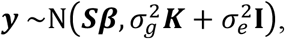

with ***S***: = [**1, *X***_*i*_,***X***_*j*_, ***X**_i_**X**_j_* or ***S***: = [***1,X***_*i*_] denoting the design matrix with the locus or epistatic interaction of interest. The bacterial sequences are first binary-encoded in ***X*** with ***X**_t_* = 0 as major and ***X***_*i*_ = 1 as minor allele, and then standardized for the purpose of association testing. 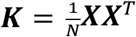 represents the kernel matrix that is used to correct for population structure. Single-locus associations are tested for statistical significance using a Wald’s-test, while epistatic associations are tested using a likelihood-ratio test comparing the full interaction model to a null model disregarding the interaction effect. Bonferroni correction was used for multiple-testing correction at *α* = 0.05.

### Implementation

The undirected graphical model was inferred using PLMC (https://github.com/deb-biemarkslab/plmc). The linear mixed model was implemented in Python and is based on Pylmm^14^. The remaining data analysis was performed in Python and Jupyter notebooks.

## SUPPLEMENT

**Supplementary Table 1**. Minimum inhibitory concentration of exploratory and confirmatory strains and their NCBI SRA identifiers.

**Supplementary Table 2**. Significant evolutionarily coupled loci in the exploratory dataset.

**Supplementary Table 3**. Entire single-locus GWAS analysis of the exploratory and confirmatory dataset.

**Supplementary Table 4**. Confirmed and Bonferroni-corrected epistatic associations.

**Supplementary Table 5**. Entire epistatic GWAS analysis of the exploratory and confirmatory datasets.

**Supplementary Figure 1.**
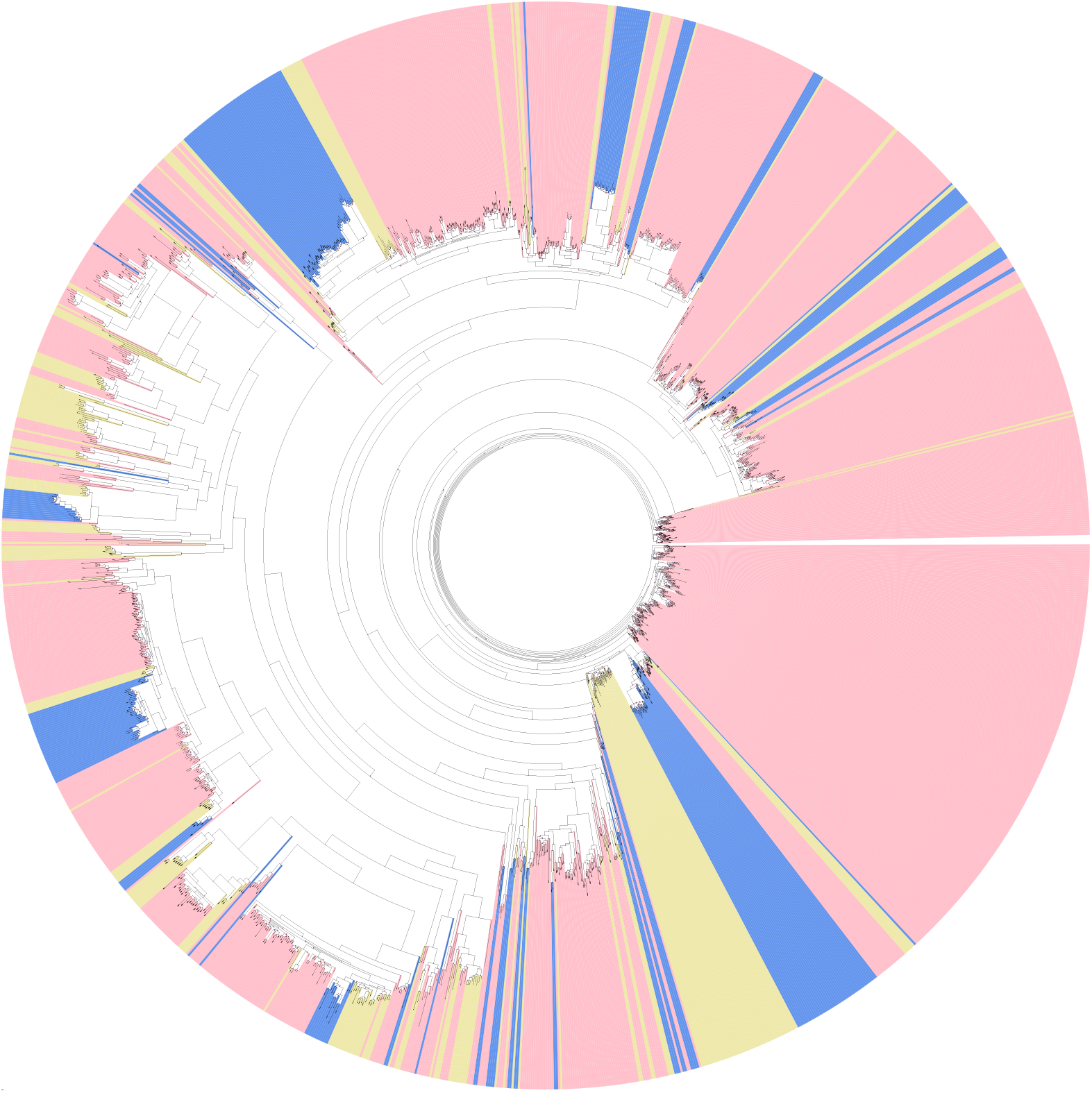
Population structure of the 1,102 exploratory (USA) and 495 confirmatory (England and Canada) *N. gonorrhoeae* isolates. USA isolates are colored pink, England isolates are yellow, and Canada isolates are blue.

